# Comparing detectability patterns of bird species in small ponds using multi-method hierarchical modelling

**DOI:** 10.1101/675116

**Authors:** José M. Zamora-Marín, Antonio Zamora-López, José F. Calvo, Francisco J. Oliva-Paterna

## Abstract

Robust knowledge of biodiversity distribution is essential for designing and developing effective conservation actions. However, monitoring programmes have historically assumed all species are detected equally with no spatial or temporal differences in their detection rates. However, recently, interest in accounting for imperfect detection has greatly increased in studies on animal diversity. In this respect, birds are the most widely used group for hierarchical occupancy-detection modelling, mainly due to the relative ease of sampling and the large number of bird datasets that are available. Nevertheless, there are no studies that have tried to evaluate the effectiveness of different bird sampling methods based on a hierarchical modelling approach. In an attempt to remedy this situation, we conducted point transects (PT), point transects plus video monitoring (PV) and mist netting (MN) in 19 small ponds located in the province of Murcia, southeastern Spain, one of the most arid regions of Europe. Multi-method hierarchical models were fitted to the detection histories of 36 common bird species with three main objectives: to compare the effectiveness of the three sampling methods for detecting the bird species using ponds, to assess the effect of sampling date on species detectability, and to establish the influence of body size and diet on species detectability. The results showed PV to be the most effective sampling method for detecting species occupancy, although detection rates ranged widely among bird groups, and some large species were weakly detected by all the methods. Average detectability increased during the breeding period, a pattern shown similarly by all sampling methods. Our approach is particularly applicable to both single- and multi-species bird monitoring programmes. We recommend evaluating the cost-effectiveness of available methods for sampling design in order to reduce costs and improve effectiveness.

## Introduction

Monitoring biodiversity is key to providing measures of status and trends of wildlife as well as for understanding its responses to threats derived from human activities. Abundance and distribution are the most widely used biological measurements in ecological studies and are frequently provided by large-scale monitoring programmes [1]. However, despite their importance for biodiversity management and conservation, most programmes are under-resourced [2], placing constraints on the number of target species, sampling effort and kind of sampling methods used to detect the target species [3]. Such limitations in survey designs may well contribute to large biases in detection probabilities, leading to the misinterpreting of abundance and distribution estimates. Indeed, concern about bias in species detectability has historically been expressed by ecologists [4], but these concerns have only begun to be explored from an analytic point of view in the two last decades. Recently, some studies have reported strongly inaccurate abundance estimates as a result of not taking into account possible imperfect detection, masking trends and providing misinformation that can affect conservation actions [5,6]. Hence, setting an accurate study design based on effective sampling methods that maximize species detectability is key to outlining any biological monitoring programmes.

The detection rate or detectability (*p*) is defined as the probability of detecting at least one individual of a given species in a particular site, given that individuals of that species are present in the study area during the survey [7,8]. Traditionally, the vast majority of studies have assumed all species composing a biological community are similarly detected [4],and detectability is constant over space, time, different methods or weather conditions. However, interest in incorporating imperfect detection (*p* < 1) into biodiversity studies has largely increased in the last two decades due to the development of novel hierarchical modelling techniques [9,10]. Several approaches have been suggested in this modelling framework in order to estimate species richness, abundance and distribution corrected for incomplete detection. For example, species richness models enable unbiased estimates of site-specific species richness to be calculated while accounting for imperfect detection [11], thus enhancing richness predictions in studies that tended only to use observed richness [6]. On the other hand, binomial *N*-mixture models allow the species abundance to be calculated with data from unmarked individuals [12], thus estimating the detection probability of a single individual from a particular species [13].However, it should be noted that some estimation failures have recently been reported for binomial *N*-mixture models and its statistical design has not been resolved [14]. Lastly, species distribution models (also known as *occupancy models*) can be applied to data of presence-absence counts in order to make mapped predictions or to know specific-species detectability [15]. Furthermore, many of these models also allow the incorporation of covariate relations in order to explore the influence of biotic and abiotic features on the abundance, richness or distribution of target species.

Distribution hierarchical models accounting for imperfect detection are composed of two different processes: an ecological process governed by the probability of occupancy and another observation process that is governed by the probability of detection [15,16]. The former is defined by the habitat requirements of each species and involves both the presence and distribution of target species in the study area (i.e. whether the species is or is not present). The latter process depends directly on occupancy and is governed by the same drivers (i.e. whether the target species is or is not detected). A species can only be detected in a sampling unit survey when that species is occupying the study unit. Besides drivers of occupancy, the observation process is constrained by several additional factors that hinder or modulate the detectability of species. These factors are derived firstly from specific-species traits, such as behaviour, life history and phylogenetic relatedness [17,18], and secondly from study design features, such as time of survey [19], sampling method, survey effort (number of surveys and sampling units), weather conditions, surveyor skills and habitat characteristics among others [15,18]. Presence-absence data across several surveys of the sampling units are required to estimate the probability of detection for any species. However, some different extensions have recently been applied to single-visit datasets in order to deal with this constraint; for example, it is possible to account for multiple independent observers, multiple independent detection methods (multi-method) or by the spatial subsampling of the study area [15,20].

Currently, birds are the most frequently used group for hierarchical occupancy-detection modelling, probably due to the greater number of datasets and statistical methods available [4]. To date, most bird studies have accounted for imperfect detection by using data from visual and aural point counts [17,19,21,22]. However, a similar effectiveness for detecting species richness has been reported for mist netting [23,24], a sampling method based on trapping birds with nets in order to mark them individually, a technique that has been increasing over recent decades [25]. There is a large literature contrasting both sampling methodologies from descriptive approaches in terms of richness and abundance [26–29]. For example, Rappole et al. [23] used data from point counts and mist netting in tropical habitats to show different method-specific biases, which are complementary to each other, and they proposed a combined methodology to provide a more accurate assessment of the avian community. Despite similar effectiveness in detecting species richness, most of these studies have pointed to the greater bias of mist netting for recording the abundance of bird species [24,30].

On the other hand, the rapid development of new technologies is revolutionizing biodiversity monitoring, and several devices can now be used to record large amounts of field data [2]. For example, video cameras have recently been used to explore drinking patterns of desert birds in small manmade ponds in areas of Arizona and Kalahari [31,32]. In arid and semi-arid regions, artificial water bodies such as drinking troughs and cattle ponds may represent the only drinking water sources for ensuring terrestrial biodiversity [31], thus providing a key service for wildlife. Therefore, these aquatic systems act as an ideal model habitat for detecting biodiversity and exploring detectability patterns in areas with scarce water availability.

To our knowledge, only three studies have aimed to explore the effectiveness of different sampling methods through a multi-method hierarchical approach, and all have mainly focused on mammal species [9,33,34], while no studies have been conducted to assess bias arising from different bird sampling methods. Here, we fit a multi-method hierarchical detection model, described by Nichols et al. [9], to contrast the effectiveness of three sampling methods for detecting 36 breeding bird species. For that purpose, 19 isolated small ponds located in a semi-arid region were selected as model habitat for the three sampling techniques to be applied. Detectability estimates were calculated for each method at species level. Furthermore, family-level effects were considered, and bird species were also grouped according to two life-history traits (body size and diet) to explore group-level effects, because phylogenetic relatedness and biological traits have been suggested as an important drivers in detectability [17,35]. Our specific aims were to: 1) compare the detection effectiveness of different sampling methods for breeding bird species; 2) assess the contribution of sampling date as a source of variation in detection probabilities during the breeding season and; 3) elucidate the influence of phylogenetic relatedness and life-history traits on detectability at method level. The occupancy-detection modelling carried out could be used as a starting point in the design stage of biological monitoring programmes, allowing resource optimization and maximizing the detectability of target species.

## Materials and methods

### Study area

This study was carried out in the province of Murcia (Región de Murcia), a province located in the southeast of the Iberian Peninsula. The study area covers 11317 km^2^, and is one of the most arid zones in continental Europe [36]. Current annual precipitation is normally less than 350 mm in most of the Iberian southeast and this ecogeographical area is characterized by a negative water balance lasting at least 4 or 5 months. Rainfall occurs predominantly in autumn and early winter, with a second short peak in early spring. In sum, the study area is characterized by a strong water deficit during spring and summer. Despite its hydrological stress conditions, the study area comprises a varied set of environments that differ in climate, topography and vegetation. In general, the inland zones have a more extreme, continental climate with colder winters and higher mean annual precipitation than the coastal zones. Generally, the Iberian southeast is occupied by mosaics of agricultural and forest areas with different degrees of representativeness. Human activities, especially agriculture, have had profound effects on the landscape structure of these environments, ranging from increased landscape heterogeneity to severe land degradation. The clear land use segregation can be explained by terrain roughness and human occupancy. Dry and irrigation farming predominate in the lowlands and highland steppes, whereas steeper areas are occupied by Mediterranean low shrubland and pine forests (mainly *Pinus halepensis*).

During recent decades, land uses in this area have been increasingly devoted to extensive agricultural irrigation practices, which, together with the natural water scarcity, have led to the overexploitation of groundwater and surface water resources. This situation has dramatically decreased the free water available to wildlife [37], especially in seasons of water deficit. Thus, the small isolated ponds still present in the study area, such as drinking troughs and artificial pools, play an essential role in supporting biodiversity [38–40] and act as shelters for animal species linked to aquatic ecosystems [41].

Ponds provide several key services to terrestrial fauna such as surface water and food resources [42,43]. Therefore, these aquatic ecosystems have become useful model habitats in biodiversity studies due to their attraction for terrestrial animal species. In the present study, 19 small ponds extending across an inland-coastal gradient (S1 Fig), and located in predominantly agro-forestry areas, were selected. The main criteria for selecting the water bodies studied were: (1) good access conditions for drinking terrestrial birds and their regular use by the avian community, and (2) the absence of pond features (surrounding habitat, vegetation cover, availability for birds, etc.) that would affect detectability. As a whole, the selected sampling sites are mainly used for cattle and game-species watering or for storing water. Although most of the study ponds are fed from natural springs, all of them have been slightly modified by lining them with cement to ensure permanent water availability.

### Sampling design

We used detection-non detection data from point transects (PT), video cameras monitoring and mist netting (MN) conducted around the 19 study ponds. Sites were sampled three times with every sampling method from 28 March to 28 July 2017 (sampling period), covering the breeding season of birds in the study area. The three surveys for each sampling site were conducted in early-mid spring, late spring and early summer, respectively. These sampling methods were successively applied at the study ponds, where PT and video camera monitoring were the first to be applied to avoid possible behavioural changes in the birds caused by the more invasive method of MN [44]. PT were carried out in a portable hide deployed on the vegetation surrounding the ponds, where it was not expected to influence bird activity. The hide was 10 m from the pond and binoculars were used for species identification. All birds seen or heard in or around the study ponds (up to 10 m) were recorded. Conventional video cameras were used as a complement to PT, this combination between PT and video cameras being termed as point transects and video monitoring (PV). Conventional video cameras (Panasonic Handycam, HC-V180, Panasonic Corporation, Osaka, Japan) were deployed in 10 sites, where an additional small pool (filling from the main pond) was out of the watcher’s view, in order to record all the birds using the pool. Cameras were positioned to cover the entire surface of the pools to ensure the birds were detected when drinking at any edge of the water. The use of video cameras as effective sampling method to record birds drinking at ponds has recently been reported [31]. Videos were later analysed in the laboratory by visualizing the entire video records, and similar data as PT were recorded. As a result, two different visual methods could be described: PT, which accounts for bird records only detected by the surveyor, and PV, which accounts for bird records jointly detected by surveyor and camera.

MN seasons were based on three nets (2 x 12 m and 1 x 9 m, 16 mm mesh size) open in a 10 m radius round the ponds and deployed between the water and surrounding vegetation to decrease net visibility. Once captured each bird was ringed, measured (data not used in this study) and released. MN was conducted in eight ponds where conditions were suitable to open the nets; for example, nets could not be opened in sites with high density vegetation surrounding the pond.

Periods between surveys at each site were no longer than 40 days and the survey order remained unchanged during the sampling period. In the study area, bird species of the coastal region showed a slightly advanced breeding phenology due to warmer conditions. Thus, littoral ponds were the first sites to be surveyed in order to correct for this phenomenon and last ones were the ponds located in wetter inland areas. Each sampling lasted 3 hours, beginning at sunrise and in good weather conditions [42]. The early morning period has been described as the time of greatest bird activity, after which species detectability steeply declines [26,29]. As mentioned, the three sampling methods were applied in similar conditions (sampling effort, good weather conditions and the same sampling range time), and so it is assumed that they provide representative information about bird community during the sampling periods, while any difference in the results can be attributed to methodology biases [30].

### Detection histories

We generated method- and survey-specific detection histories for all the breeding species recorded during the study period. Species with less than five records were removed from the models in order to avoid bias related to small sample size, since estimates may be unreliable when information collected on species presence/absence is limited [45,46]. Additionally, migratory non-breeding species in the study area were also removed from the models to meet the closure assumption. Separate detection histories were generated for each sampling method and each survey visit. Therefore, as sampling event refers to one survey per method, a maximum of nine detection events were possible for each species because three surveys were conducted for each of the three sampling methods. A value of 1 was attributed when a species was detected in a specific survey using a single method, a value of 0 being given otherwise. Thus, for a given species, the detection history of 010 101 000 meant one or more individuals were detected only at the second survey by the first sampling method, the presence of the same species was recorded at the first and third surveys by the second method, and no species presence was detected by the last sampling method.

### Modelling framework

Multi-method hierarchical occupancy modelling, as described by Nichols et al. [9], an extension of other occupancy models [45], was used to estimate the detectability of bird species recorded during the survey period. Using this approach, method-specific detection probabilities can be calculated for two or more sampling methods [9,33]. The detection process is modelled as a Bernoulli random variable as follows:

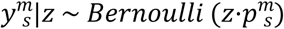

where 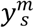 represents detection/non-detection records for method *m* and survey *s, z* denotes occupancy status (0/1), which is constant over time, and 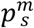 is the probability of detection using method *m* and survey *s*.

All analyses were carried out using a maximum likelihood-based approach implemented in the Presence version 10.6 programme [47] through R interface (version 3.4.3.). Four models were fitted to account for the variability derived from interference of sampling methods and survey occasions on species detectability (Table 1): a null model (*p*), assuming probability of detection as constant for all sampling methods and surveys; a method-specific model (*p^m^*), assuming detectability as dependent on methods but constant between surveys; a survey-specific model (*p_s_*), as time-dependent detectability but constant between methods; and method- and survey-specific model 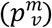, with surveys as an additive effect with methods. The species-specific occupancy probability (*ψ*) and site-specific occupancy probability (*θ*) were assumed to be constant over time and for the whole study area.

**Table 1.** Multi-method models fitted to estimate detectability of bird species in ponds located in the province of Murcia. In all cases the occupancy probability (*ψ*) and specific-site occupancy probability (*θ*) were modelled as constants. The probabilities of detection were modelled as constant (*p*), as specific of the method (*p^m^*), as specific of the survey (*p_s_*), and depending on both method and survey 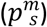.

| Name | Model | Structure (detection component) |
| --- | --- | --- |
| Null | $\psi, \theta, p$ | $\text{logit}(p) = b_0$ |
| Method | $\psi, \theta, p^m$ | $\text{logit}(p^m) = b_0 + b_m$ |
| Survey | $\psi, \theta, p_s$ | $\text{logit}(p_s) = b_0 + b_s$ |
| Method+survey | $\psi, \theta, p^m_s$ | $\text{logit}(p^m_s) = b_0 + b_m + b_s$ |

$b_m$ , covariate specific of method; $b_s$ , covariate specific of survey.

Model-averaging of the four hierarchical models was followed to calculate the averaged estimates of occupancy probability and detection probability at species level. The normalized weights of Akaikés Information Criterion (AIC_w_) were used to establish the weight of evidence in support of each model. Differences in AIC (ΔAIC) between each model and the best one were used to rank models [48,49] and establish the overall importance of both variables (sampling method and survey date) in explaining our estimates.

The sampling design considered the analytical assumptions required to fit hierarchical occupancy models [1,50]. We assumed that the communities studied were closed during the sampling season. The survey period lasted four months, from 28 March to 28 July, overlapping with the breeding season of all the terrestrial bird species of the study area. During this time, most breeding species are settled in their breeding territories and large movements are not expected. However, the spring migration of some species overlapped with our study period and migrant species were detected on the first survey occasion at each pond. To meet the closure assumption, all migratory non-breeding species in the study area were removed from the modelling. On the other hand, weather conditions and sampling effort (detection time) were similar for all surveys so were not expected to affect detectability. We also assumed that occupancy was independent among study sites because the minimum distance between ponds was always greater than 1.5 km, which is a reasonable distance to consider sites as independent when the survey period covers the breeding season of birds.

### Species groups

Phylogenetic relatedness and biological traits of the recorded species were used to explore detectability patterns at family level and at group level in order to assess bias related to morphological, phylogenetic and trophic variables. Recently, phylogeny has been suggested as a driver of detectability in bird species, since it affects singing behaviour, thus influencing detection probability [17,51]. However, some recent studies suggest that life-history traits may also affect species detectability [17,52]. Among these traits, body size (morphological variable) and diet (trophic variable) have previously been used in hierarchical occupancy modelling [46] and both are expected to influence detectability, particularly in our study case. We used both biological traits to allocate species to bird groups. Body mass was used as a measure of body size [30,46], because it is a reasonable indicator of bird total size. Thus, bird species were grouped into three body size classes (small, body mass <30 g; medium-sized, body mass 30-100 g; large, body mass >100 g) and four trophic classes were established according to diet (insectivorous, seed-eaters, frugivorous and generalists). Life-history traits of the studied species were obtained from Pearman et al. 2014 [53]. Taking into account both life-history traits, we obtained six bird groups based on information about size and diet of the recorded species: (1) small insectivorous, (2) medium-sized and large insectivorous, (3) small frugivorous and insectivorous, (4) small seed-eaters, (5) medium-sized and large seed-eaters, and (6) medium-sized and large generalists.

### Ethics statement and permits

Authorization for our study was provided by the Dirección General de Medio Ambiente of the Autonomous Community of Murcia (reference number: AUF20170002), which regulates the management of wildlife in the study area. Ringing license was provided by the Spanish Ministry of Agriculture, Fisheries and Environment. Most of the study ponds were located on public land, while permission for access was sought from landowners in the case of ponds located on private land.

## Results

A total of 5304 birds belonging to 36 species recorded in small ponds during the sampling period were used for modelling occupancy and detectability (Table 2). Another 26 taxa belonging to migratory non-breeding birds in the study area, such as the Pied Flycatcher and Willow Warbler, and occasional species with less than 5 records were removed from the statistical analysis. The results from model averaging pointed to the survey-specific model as the best supported model for 41.7 % of bird species (15 taxa) based on AIC, closely followed by the method-specific model and null model for 33.3 % (12 taxa) and 25 % (9 taxa)of the species, respectively (Fig 1, S1 Table). The model with survey as an additive effect was largely unsupported because it was not the best for any species.

**Fig 1.**
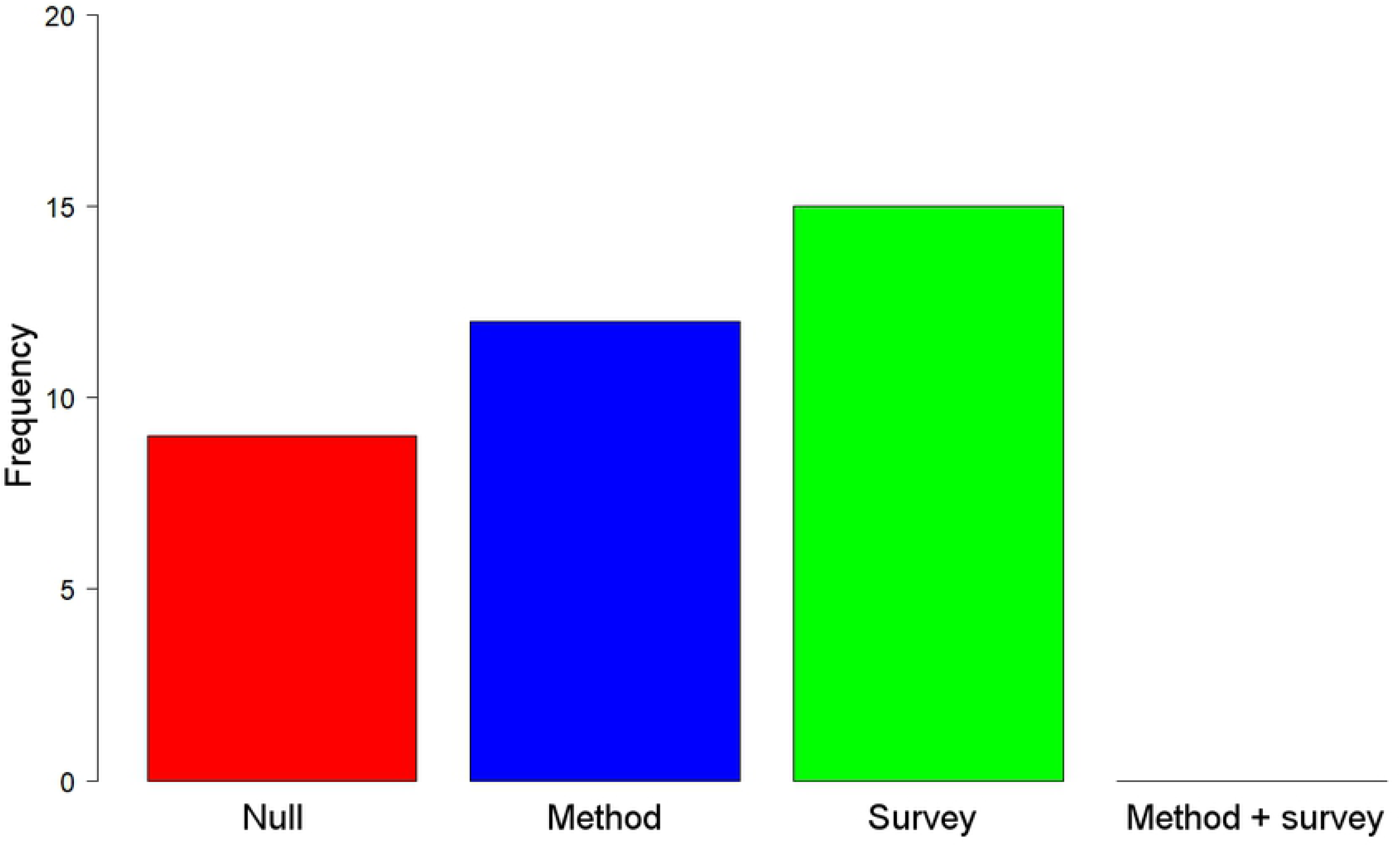
Frequency distribution of best models explaining detection estimates of bird species recorded on ponds in the province of Murcia. The selection procedure was based on the lowest AIC score. Null model assumes detection probability as constant across methods and surveys; method-specific model assumes detectability as dependent on methods but constant over the surveys; survey-specific model considers time-dependent detectability but constant for all methods; and method- and survey-specific model assumes detectability as dependent on surveys as an additive effect with methods.

**Table 2.** Summary of species name, family and group membership for breeding bird species recorded during pond surveys in the province of Murcia. Another 26 bird species were recorded in pond surveys but they were removed from modelling due to their non-breeding status in the study area or their small sample size.

| Species | Common name | Family | Bird group<sup>a</sup> |
| --- | --- | --- | --- |
| *Aegithalos caudatus* | Long-tailed Tit | Aegithalidae | 1 |
| *Carduelis cannabina* | Common Linnet | Fringillidae | 4 |
| *Carduelis carduelis* | European Goldfinch | Fringillidae | 4 |
| *Carduelis chloris* | European Greenfinch | Fringillidae | 3 |
| *Certhia brachydactyla* | Short-toed Treecreeper | Certhiidae | 1 |
| *Columba palumbus* | Common Woodpigeon | Columbidae | 5 |
| *Cyanistes caeruleus* | Eurasian Blue Tit | Paridae | 1 |
| *Emberiza calandra* | Corn Bunting | Emberizidae | 5 |
| *Emberiza cia* | Rock Bunting | Emberizidae | 4 |
| *Emberiza cirlus* | Cirl Bunting | Emberizidae | 4 |
| *Erithacus rubecula* | European Robin | Muscicapidae | 1 |
| *Fringilla coelebs* | Common Chaffinch | Fringillidae | 4 |
| *Garrulus glandarius* | Eurasian Jay | Corvidae | 6 |
| *Hippolais polyglotta* | Melodius Warbler | Acrocephalidae | 3 |
| *Lanius senator* | Woodchat Shrike | Laniidae | 2 |
| *Lophophanes cristatus* | Crested Tit | Paridae | 1 |
| *Loxia curvirostra* | Red Crossbill | Fringillidae | 5 |
| *Luscinia megarhynchos* | Common Nightingale | Muscicapidae | 3 |
| *Muscicapa striata* | Spotted Flycatcher | Muscicapidae | 3 |
| *Parus major* | Great Tit | Paridae | 1 |
| *Periparus ater* | Coal Tit | Paridae | 1 |
| *Petronia petronia* | Rock Sparrow | Passeridae | 6 |
| *Phoenicurus ochruros* | Black Redstart | Muscicapidae | 1 |
| *Phylloscopus bonelli* | Western Bonelli's Warbler | Phylloscopidae | 1 |
| *Phylloscopus collybita* | Common Chiffchaff | Phylloscopidae | 1 |
| *Pica pica* | Eurasian Magpie | Corvidae | 6 |
| *Saxicola torquata* | Common Stonechat | Muscicapidae | 1 |
| *Serinus serinus* | European Serin | Fringillidae | 4 |
| *Sitta europaea* | Eurasian Nuthatch | Sittidae | 3 |
| *Streptopelia turtur* | European Turtle-dove | Columbidae | 5 |
| *Sylvia cantillans* | Subalpine Warbler | Sylviidae | 3 |
| *Sylvia hortensis* | Western Orphean Warbler | Sylviidae | 3 |
| *Sylvia melanocephala* | Sardinian Warbler | Sylviidae | 3 |
| *Sylvia undata* | Dartford Warbler | Sylviidae | 3 |
| *Turdus merula* | Eurasian Blackbird | Turdidae | 2 |
| *Turdus viscivorus* | Mistle Thrush | Turdidae | 2 |

<sup>a</sup> Bird groups are based on body size and diet traits. Numbers refer to six different groups: 1, small insectivorous (<30g); 2, medium-sized and large insectivorous (≥30g); 3, small insectivorous and frugivorous (<30g); 4, small seed-eaters (<30g); 5, medium-sized and large seed-eaters (≥30g); and 6, medium-sized and large generalists (≥30g).

Model-averaging estimates of detection probability for study species showed clear differences in method-specific detectability (Fig 2). Occupancy detection increased slightly during breeding season but differences among the three sampling methods remained constant for the three surveys. PT and PV provided similar detectability estimates but slightly higher in the second case. PV obtained detectability estimates significantly higher than MN, but the other pairwise comparisons did not provide significant differences. Nevertheless, detection estimates of some species were low even for PV. MN provided the lowest detectability estimates of the three studied methods. Moreover, MN showed the highest variability in species detectability because this method covered a wide range from almost full detection (*p ≈ 1*) for some species (e.g. European Greenfinch and European Serin) to practically null detection (*p ≈* 0) for others such as Common Woodpigeon.

**Fig 2.**
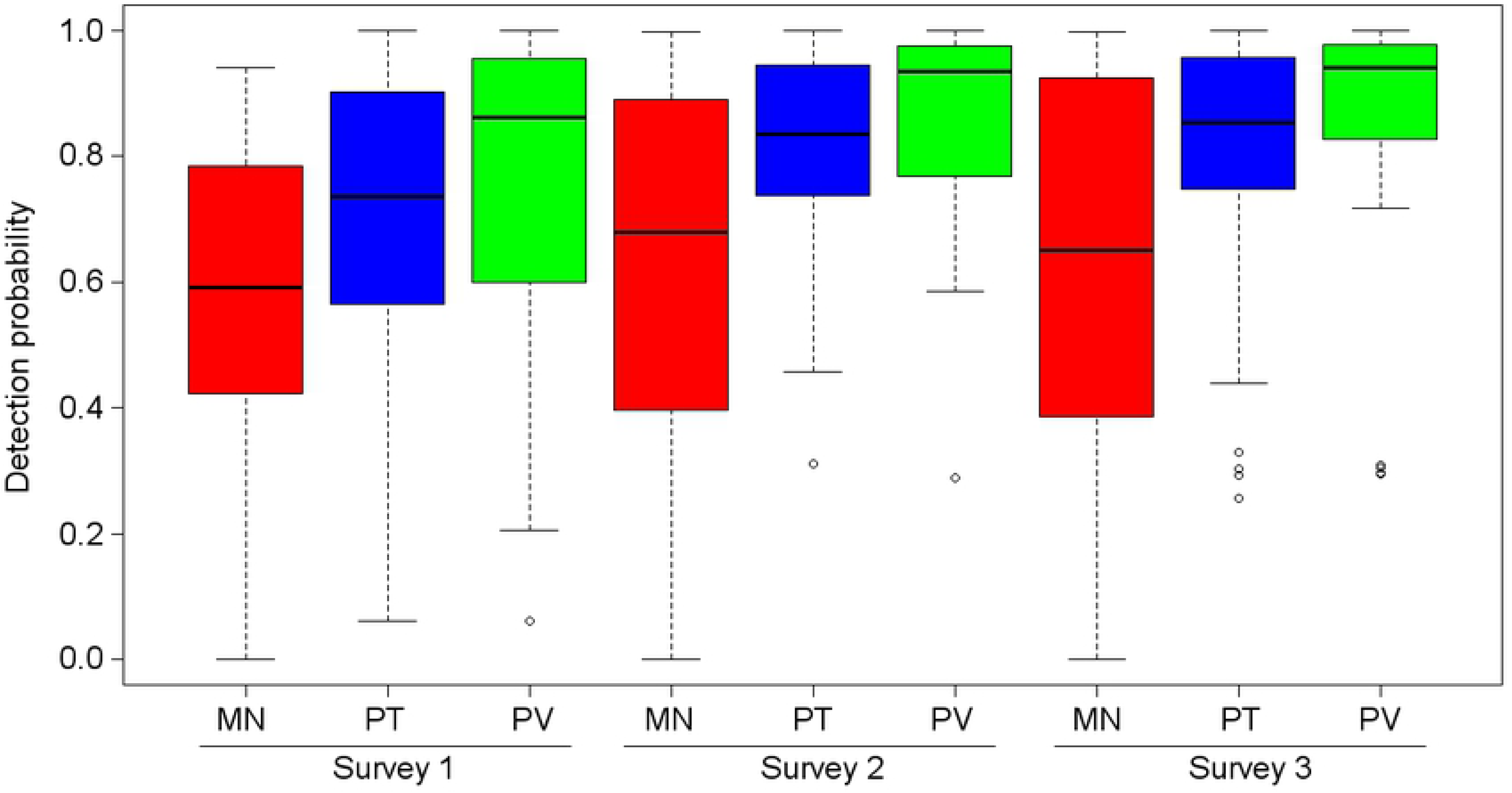
Boxplot of the detection estimates of bird species based on model averaging for each sampling method and survey occasion. The three sampling methods are indicated: MN, mist netting; PT, point transects; and PV, point tranects plus video monitoring. Surveys 1, 2 and 3 correspond to visits made in early-mid spring, late spring and early summer, respectively.

No relation between species occupancy and species detectability averaged between sampling methods was found (Fig 3A). The occupancy estimates ranged widely from *ψ* = 0.14 in Eurasian Nuthatch to *ψ* = 1 in Eurasian Blackbird. However, the detection estimates for many studied species (89.9%) was higher than *p* = 0.5, and only four of the 36 modelled species showed lower values. It should be noted that two of these four species, Common Chiffchaff and Corn Bunting, exhibited a relatively low detectability in spite of the fact that their probability of occupancy was complete (*ψ* = 1; *p < 0.5*0). Finches were the family with the highest occupancy and detection estimates, all finch species showing *ψ* > 0.80 and *p* > 0.80, with the exception of European Greenfinch, which exhibited a low estimated occupancy value (*ψ* = 0.35; *p =* 0.94).

**Fig 3.**
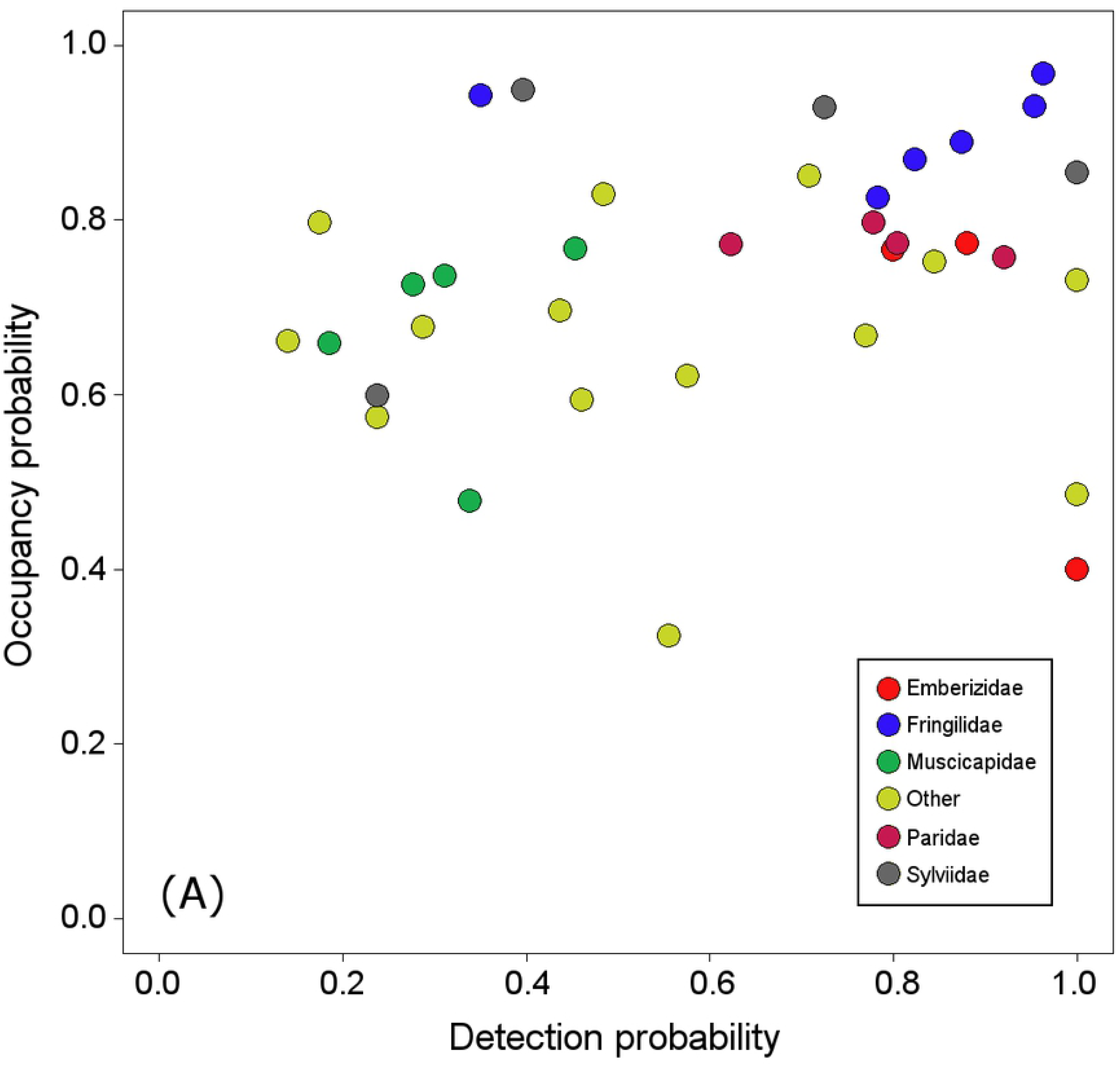
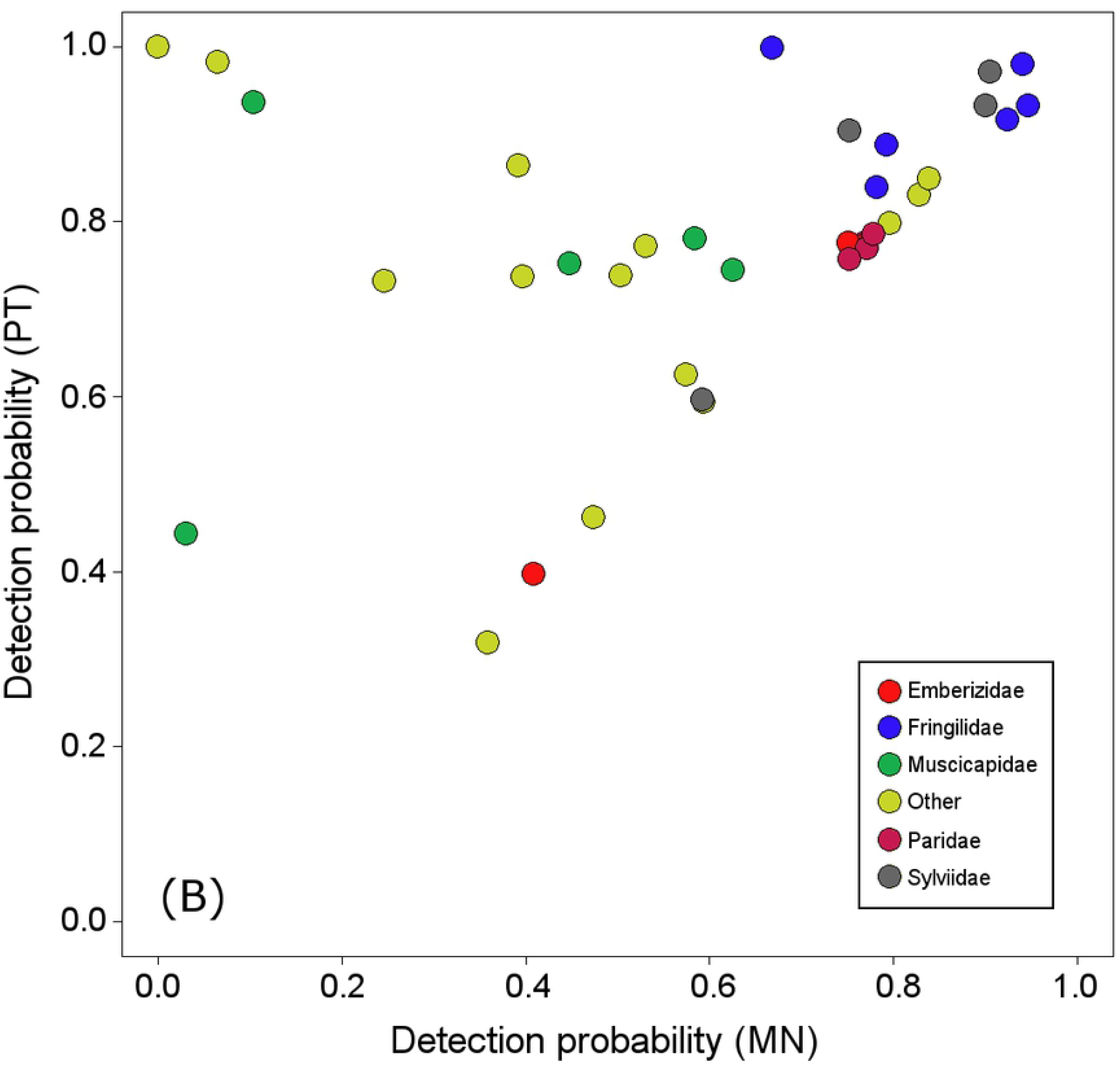
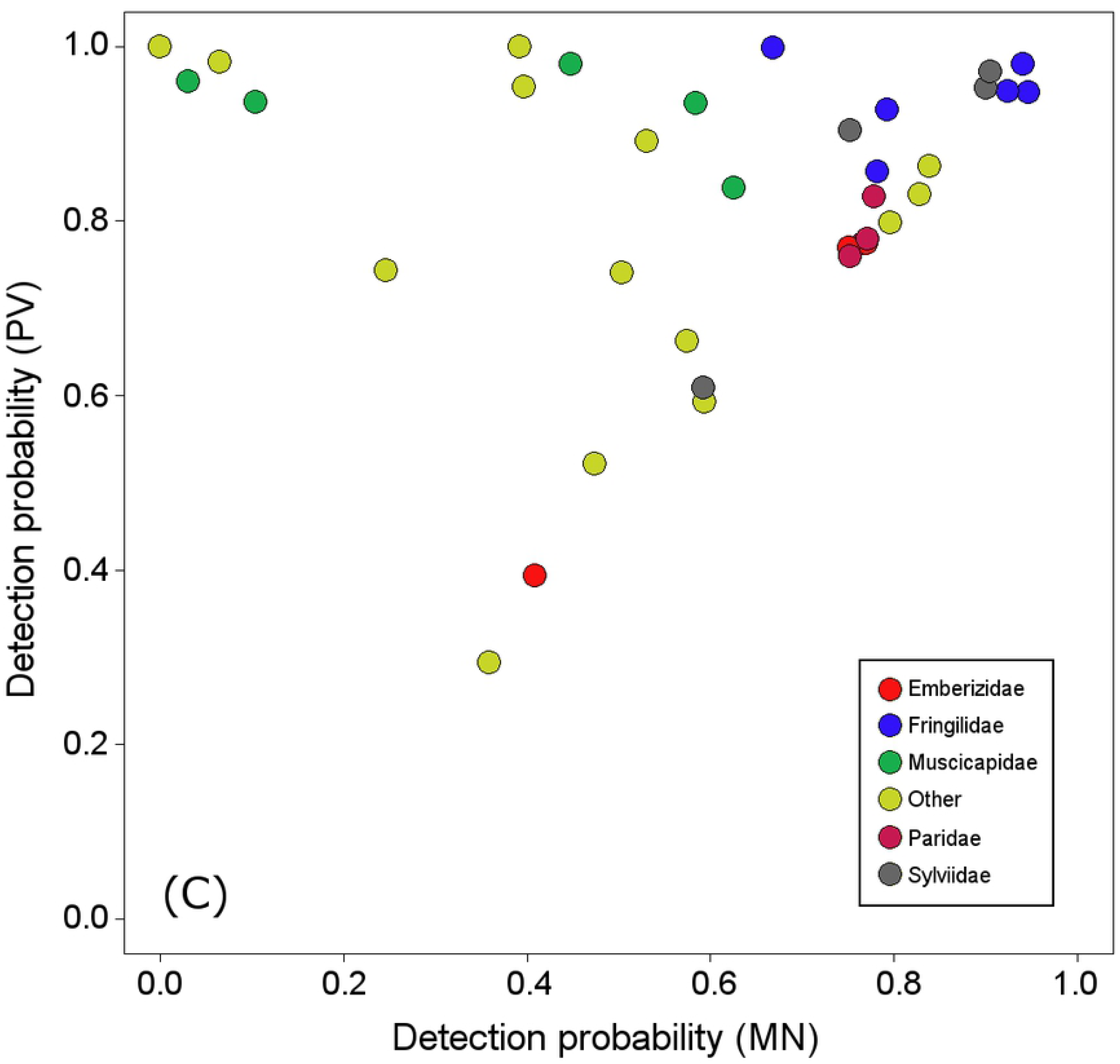
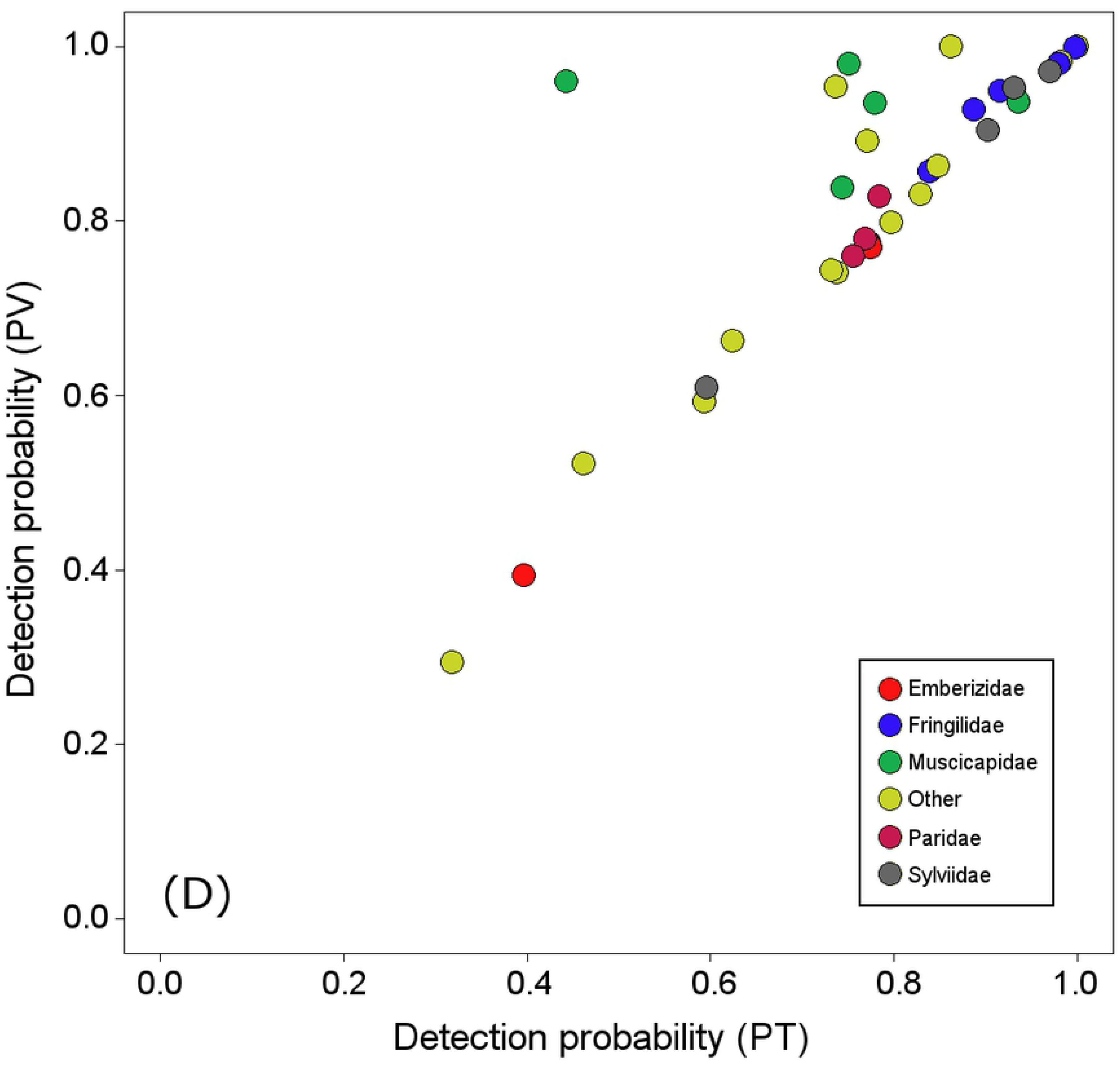
Occupancy estimates and detection estimates of bird species recorded on ponds in the province of Murcia. Tag color refers to avian family. Only the five families with the highest number of recorded species are indicated, while the remaining families are grouped as Other. (A) Occupancy probability *versus* detection probability, the latter averaged for the three target methods. (B) Pairwise comparison of detection estimates from MN (mist netting) and PT (point transects). (C) Pairwise comparison of detection estimates from MN (mist netting) and PV (point transects plus video monitoring). (D) Pairwise comparison of detection estimates from PT (point transects) and PV (point transects plus video monitoring).

A lineal relation was clearly seen for the estimated detectability using each method when pairwise comparisons of sampling methods were carried out (Fig 3B-3D). Of the 36 bird species recorded, 18 were equally detected by the three methods since their estimates at method-level differed by less than 5% among them, indicating effectiveness of these sampling methods was very similar for detect these species. However, both observational methods (PT and PV) were much more effective than MN in detecting bird species such as Common Woodpigeon, Eurasian Magpie, Spotted Flycatcher, European Turtle-dove and Eurasian Jay. Otherwise, MN was not more effective than observational methods for any species modelled, except Short-toed Treecreeper whose estimate was very slightly higher with MN (*p_PT_* = 0.31; *p_MN_* = 0.35).

Six of the 36 modelled species were only recorded by observational methods, which corresponded to large birds (such as European Turtle-dove, Eurasian Magpie and Common Woodpigeon) or species with few records (*n* < 10, such as Common Nightingale and Woodchat Shrike). However, no species were detected by MN alone. Contrasting results at family level were found in the case of method-specific detectability. Observational methods showed significantly higher effectiveness than MN for detecting the Muscicapidae family (flycatchers) and remaining families grouped as Other, and a slightly higher effectiveness for the Fringillidae and Sylvidae families (finches and warblers, respectively). However, the estimated detectability of Emberizidae (buntings) and Paridae (tits) families was very similar for the three sampling methods (Fig 4). Detectability with the PT and PV methods was very similar for all the studied families, except Muscicapidae family which were significantly better detected by PV. On the other hand, visual methods were also more effective than MN at detecting species at group-level (Fig 5). Only small seed-eaters were equally detected by the three sampling methods. Detectability for the other groups was always higher with PT and PV than with MN. Detection probability for insectivores and frugivores species (groups 1-3) showed a similar pattern, increasing slightly from MN to PT and PV. Moreover, visual methods were significantly more effective than MN at detecting medium-sized and large see-eaters and generalists. Detectability by PT and PV was very similar for all bird groups.

**Fig4.**
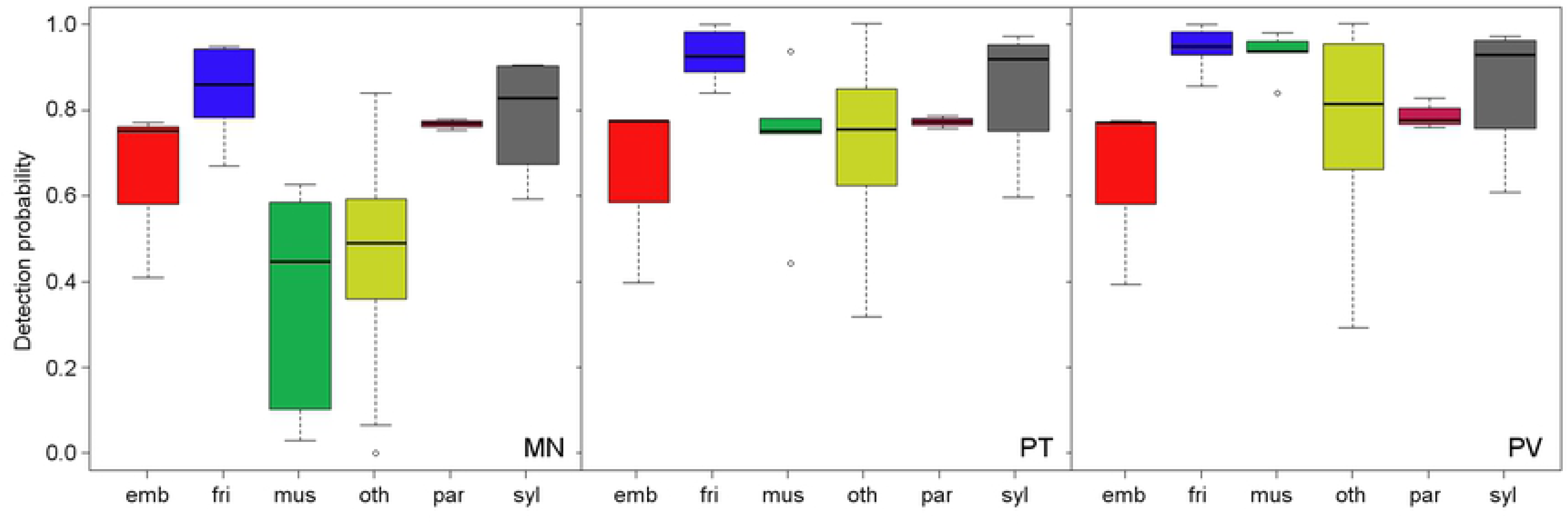
Estimated detectability at family level for each of the three sampling methods deployed in Murcia to record bird species using ponds. Sampling method label appears in the bottom right corner as follows: MN, mist netting; PT, point transects; and PV, point transects plus video monitoring. Only the five families with the highest number of recorded species are indicated, while the remaining families are grouped as other. Families are indicated as follows: emb, Emberizidae; fri, Fringillidae; mus, Muscicapidae; oth, Other; par, Paridae; and syl, Sylvidae. Surveys 1, 2 and 3 corresponding to visits conducted in early-mid spring, late spring and early summer, respectively. Box color is shown for a better understanding.

**Fig 5.**
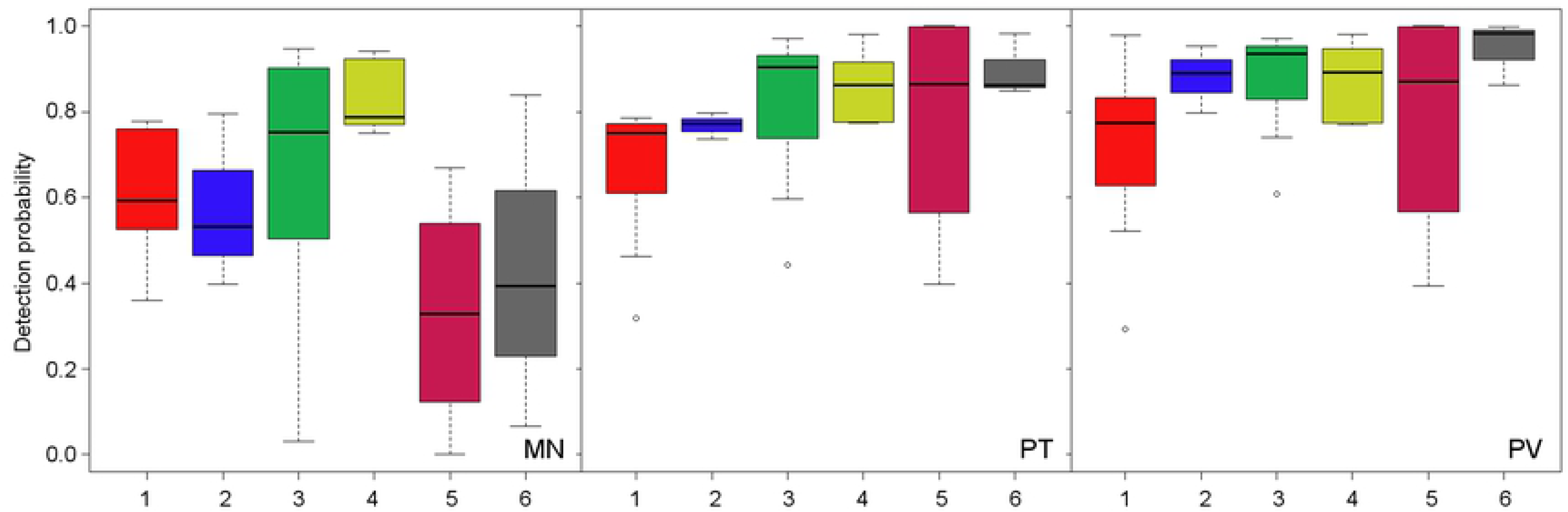
Estimated detectability at group level for each of the three sampling methods deployed in the province of Murcia to record bird species using ponds. Sampling method label appears in the bottom right corner as follows: MN, mist netting; PT, point transects; and PV, point transects plus video monitoring. Bird groups are based on body size and diet traits. Numbers refer to six different groups: 1, small insectivorous (<30g); 2, medium-sized and large insectivorous (≥30g); 3, small insectivorous and frugivorous (<30g); 4, small seed-eaters (<30g): 5, medium-sized and large seed-eaters (≥30g); and 6, medium-sized and large generalists (≥30g). Surveys 1, 2 and 3 corresponding to visits conducted in early-mid spring, late spring and early summer, respectively. Box color is shown for a better understanding.

Detectability over the whole survey period was very similar for almost all the avian families (S2 Fig). Buntings were the only family whose detectability clearly increased over time. As regards bird groups, most of them showed an increasing pattern as surveying progressed, with the highest detectability value being attained in the last survey (S3 Fig). Differences in detectability among bird groups decreased with time. Survey-specific detection estimates for each of the three sampling methods are reported on S3 Table.

## Discussion

Few studies have fitted hierarchical models focused on entire bird communities to estimate both detection and occupancy probability. In this study, we used an occupancy-detection modelling approach to assess imperfect detection in bird datasets derived from three different sampling methods. This hierarchical approach allowed us to calculate both method-specific and survey-specific detection estimates for 36 breeding bird species associated with small ponds, which represent 30.0 % of the terrestrial breeding bird community in the study area [54]. To our knowledge, this is the first attempt to compare the effectiveness of different sampling methods for bird monitoring based on an occupancy-detection modelling approach.

### Multi-method detectability

PV was significantly more effective for occupancy detection than MN, and the same method was slightly more effective than PT. Detection probability gradually increased as the breeding season progressed for the three sampling methods. Previous studies have pointed to a contrasting influence of survey date on bird detectability. For example, several species have shown unchanged detectability with time, whereas others have shown a strongly increasing or sharply decreasing time-dependent detectability [19]. A possible reason to explain our result is that we violated the closure assumption of bird species during the survey period, one of the obligatory analytical assumptions for occupancy-detection modelling [15,50]. The surveys were conducted during the dry season (spring and summer) around small ponds in a semiarid region and the drinking rates of birds may not have been constant over the whole survey period. Indeed the bird’s consumption of water probably increased with increasing temperature toward the latter half of the survey period when the days were hotter. In arid zones, birds have been reported to use water bodies more frequently during hot periods [31,55], species detectability increasing as a consequence.

Unsurprisingly the detection estimates for both visual methods were very similar, suggesting the additional use of video cameras provides only a slight improvement over the results obtained by the most traditional method of PT. However, the video cameras substantially increased the detectability of Turdidae family and, especially, some species such as Common Nightingale (*p_PT_* = 0.44; *p_PV_* = 0.96). In this context, it should be noted that additional use of video cameras may be regarded as a useful monitoring tool in biodiversity studies focused on the pull-effect of resources, such as ponds or animal feeders [31,56], where species detectability can be greatly improved by using these devices. The average detectability for the three methods showed similar estimates for closely related species. Warblers and finches were the avian families with highest detectability, with eight species having a probability of detection ranging from 0.85 to 0.96. Accordingly, phylogenetic relatedness has been reported as a driver of species detectability so that closely related taxa are expected to show similar detection rates [17]. Moreover, six of these eight species were the most abundant birds in our study, suggesting avian abundance influences the detection process, as has been reported in previous studies [15]. On the other hand, the detectability of flycatchers showed significant differences between sampling methods, PV being the best method for recording these species, closely followed by PT. The higher effectiveness of visual techniques to detect flycatchers is probably explained by their conspicuous feeding behaviour that makes them easily detectable.

MN was ineffective at detecting both medium-sized and large seed-eaters and generalist birds, such as doves and crows, or species with a very patchy distribution, such as nightingales and flycatchers. However, MN was effective at recording the presence of two small warbler species (Spectacled Warbler and Eurasian Blackcap) that were not detected by the observational methods, but they were removed from the modelling analysis due to their small sample size. These results agree with previous studies that found PT to be more effective for detecting gregarious and large birds, such as doves and crows, and conspicuous species such as flycatchers [24,30,57], while MN was more effective for detecting secretive and cryptic species [28,58,59], such as warblers. Only small seed-eaters were detected with similar effectiveness by the three target methods. Importantly, MN showed the higher variability in the detection estimates and also led to large differences in species detectability even within families and groups (Fig 4 and Fig 5). This finding underlines the view that MN should not be used as a single method to study entire bird communities, mainly because of its bias toward medium-sized and large birds.

Both visual methods provided slightly higher detectability estimates than MN for small-sized groups,contrasting with previous studies that found MN to be more effective for detecting small-sized species [58]. A possible reason to explain this result would be the use of small ponds as study model could be masking bias arising from our sampling methods. Vegetation around the water edges of our study ponds was scarce or absent, so drinking birds are generally easily detected visually independently of their behaviour. Thus, it is possible that the body size of birds did not affect detectability when observational methods were used in our study.

### Ponds as study models

We used small ponds as model habitat to inventory the breeding bird communities settled around these aquatic systems. In semi-arid environments such as the Iberian southeast water bodies exert a strong attractive pressure for terrestrial animals and they offer an interesting chance to study biological communities. Small ponds in this area are critical habitats for sustaining biodiversity due the scarcity of free water resources available to wildlife in this semi-arid region [37]. The high proportion of bird species using our study ponds is a clear example of their contribution to biodiversity. The breeding bird community of the study area consists of around 120 species, excluding marine and wetland birds [54]. We recorded 57 breeding bird species using the studied small ponds, which represent 47.5% of the terrestrial breeding bird species in the whole study area. However, all the studied ponds were located in mountainous areas dominated by Mediterranean forest, and no ponds from steppe lands or farmlands were included in the study design. Hence, typically steppe birds, such as larks and sandgrouse, also probably use ponds located in open landscapes, so that an even higher richness of birds would be expected if all types of ponds found in Iberian southeast were surveyed. Future studies that will include ponds from open areas will improve our knowledge of the services offered by these critical habitats for the conservation of terrestrial birds. Whatever the case, we recommend the use of small ponds as a supplementary and additional tool in biological monitoring programmes in arid and semi-arid environments, since they increase the ability to collect more rigorous data. For example, the implementation of pond surveys in large monitoring programmes (such as breeding bird surveys or specific surveys focused on species of conservation concern) in semi-arid regions would complement data on species distribution and so contribute to conservation actions. The power of attraction of ponds for birds leads to a high proportion of species inhabiting in their vicinity, because they can exploit one or more of the available resources (such as water to drink or bathe in, and a source of food), making them easily detectable. Moreover, our results suggest that surveying birds in ponds decreases some of the biases related with the sampling methods used in this study and improves their effectiveness.

However, the decline of extensive farming practices and the overexploitation of groundwater related to new farming practices are increasing the rate at which small ponds are being lost in Iberian southeast [37]. Therefore, a thorough evaluation of the services provided by ponds to ecosystems functioning and to supporting biodiversity is urgently needed in order to promote the conservation of these vulnerable aquatic habitats. Although the role of ponds in the conservation of freshwater biodiversity has been studied from different taxonomic points of view, there is a wide gap in our knowledge of the importance of small water bodies to terrestrial biodiversity in general [31]. Accordingly, future research efforts should focus on the importance of these critical freshwater ecosystems for the conservation of mammal and bird terrestrial communities in arid and semi-arid environments.

### Monitoring implications

Our study points to the greater effectiveness of PV and PT compared to MN for detecting bird species. However, we recommend a rigorous evaluation of the most suitable sampling method during the design stage of any study because effectiveness will depend mainly on the study aims, the study area, the target species and the available resources. For example, PT need a high degree of skill, which must be equal for all observers if species identification is to be unequivocal [44], demanding a high level of training in areas of great avian richness. However, PT is easier and faster to conduct than MN and generally demands less material, and both human and economic resources [60], making it perhaps the most effective in terms of species detected per unit effort [25,28,29]. Moreover, visual techniques are less invasive than MN and do not interfere with bird activity [30].

On the other hand, the most novel method, PV, can increase the detection rates of given species in bird studies of pull-effect resources, such as ponds, where it is not possible to see the entire water surface, so that some species would be overlooked, leading to incomplete data. In our case, the additional use of video cameras allowed us to greatly improve the detectability of several muscicapid species (Common Nightingale, Common Stonechat, European Robin and Black Redstart) and thrush species (European Blackbird and Mistle Thrush) compared with the simpler method of PT. For some of the above species, detection rates increased by more than 20% (Common Stonechat and Mistle Trush) and even 50% (Common Nightingale) when video cameras were used as a complement to PT. The use of video cameras as a single method can reduce the sampling effort by covering several sampling sites simultaneously, but it is not always possible to cover the entire surface of the target habitat. Moreover, it should be noted that conventional cameras operate continuously and the lab time to review all recorded videos is not negligible. However, the method that involves most time and human resources is MN because at least two operators are required to reduce the time during which birds are handled. Nevertheless, MN provides an easy way to standardized sampling, decreasing surveyor bias, detecting species that are often missed using other count methods and enabling handling, thus providing individual information [25]. So, MN can provide very useful data for population management, such as breeding status, body condition or the sex-ratio of the target species [60]. For example, we obtained the first evidences of breeding by Hawfinch and Common Redstart in our study area through MN conducted around some of the ponds studied. Accordingly, MN can be equally effective as PT to detect avian richness in habitats with high density vegetation and low visibility conditions, such as reed beds. The additional and invaluable information obtained may be regarded as compensating for the increased time and effort needed [60]. Evaluating the cost-effectiveness of different sampling methods, then, is recommended to match the available resources to the study aims. Our multiple method modelling approach can be especially useful in multispecies conservation programmes, acting as a starting point to design accurate surveys accounting for incomplete detection.

## Aknowledgments

We thank to colleagues and members of Departamento de Zoología y Antropología Física and Departamento de Ecología e Hidrología of the University of Murcia for their help in fieldwork, as well as members of the ANSE Bird Ringing Group, especially Francisco A. García Castellanos. We also thank the Dirección General de Medio Ambiente of the Autonomous Community of Murcia for permission to access to protected areas.

## Supporting information

**S1 Fig. Distribution of study ponds located in the province of Murcia.** Coordinates are indicated as UTM 30S and elevation increase with the intensity of the green color.

**S2 Fig. Estimated detectability at family level for each of the three surveys conducted in the province of Murcia to record bird species using ponds.** Detection estimates are averaged from the three sampling methods. Only five families with the highest number of recorded species are indicated, the remaining families being grouped as other. Families are indicated as follows: emb, Emberizidae; fri, Fringillidae; mus, Muscicapidae; oth, Other; par, Paridae; and syl, Sylviidae. Surveys 1, 2 and 3 corresponding to visits conducted in early-mid spring, late spring and early summer, respectively. Box color is shown for a better understanding. (TIFF)

**S3 Fig. Estimated detectability at group level for each of the three survey visits conducted on region of Murcia to record bird species using ponds.** Bird groups are based on body size and diet traits. Numbers refer to six different groups: 1, small insectivorous (<30g); 2, medium-sized and large insectivorous (≥30g); 3, small insectivorous and frugivorous (<30g); 4, small seed-eaters (<30g): 5, medium-sized and large seed-eaters (≥30g); and 6, medium-sized and large generalists (≥30g). Surveys 1, 2 and 3 correspond to visits conducted in early-mid spring, late spring and early summer, respectively. Box color is shown for a better understanding. (TIFF)

**S1 Table. Occupancy estimates and detectability: best model for bird species recorded in pond surveys in the province of Murcia.** Total occupancy estimates (*ψ*) and site-specific occupancy estimates (*θ*) are indicated with confidence intervals of 95 %. Four hierarchical models were fitted to account the variability derived from the influence of sampling method and survey occasion on species detectability. The model with survey as an additive effect with methods was not the best model for any species.

**S2 Table. Model comparisons to identify covariates (method or survey) influencing detectability of 36 bird species recorded in the province of Murcia using small ponds.** Akaike Information Criterion (AIC), the relative differences in AIC (ΔAIC), the Akaike weights, the total number of estimable parameters (*K*) and deviance are indicated. Models are ordered in terms of ΔAIC. Occupancy probability (*ψ*) and site-specific occupancy probability (*θ*) were modelled as constant parameters, whereas detection probability was modelled as constant (*p*), depending on the method (*p^m^*), depending on the survey (*p_s_*), and depending on both method and survey 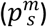.

**S3 Table. Survey-specific detection estimates from mist netting (MN), point transects (PT) and point transects plus video monitoring (PV) methods for 36 bird species recorded on ponds in the province of Murcia.** Estimated standard errors are indicated in brackets. (PDF)

## Author Contributions

**Conceptualization:** José F. Calvo, Francisco J. Oliva-Paterna, José M. Zamora-Marín

**Data curation:** José M. Zamora-Marín, José F. Calvo

**Formal analysis:** José F. Calvo, José M. Zamora Marín

**Funding adquisition:** Francisco J. Oliva-Paterna, José M. Zamora-Marín

**Investigation:** José M. Zamora-Marín, Antonio Zamora-López, Jose F. Calvo

**Methodology:** José M. Zamora-Marín, José F. Calvo, Antonio Zamora, Francisco J. Oliva-Paterna

**Project administration:** Francisco J. Oliva-Paterna, José F. Calvo, José M. Zamora-Marín

**Resources:** Francisco J. Oliva-Paterna, José F. Calvo, José M. Zamora-Marín

**Software:** José F. Calvo, José M. Zamora Marín

**Supervision:** José F. Calvo, Francisco J. Oliva-Paterna

**Validation:** José M. Zamora-Marín

**Visualization:** José F. Calvo, José M. Zamora-Marín

**Writing – original draft preparation:** José M. Zamora-Marín

**Writing – review & editing:** José M. Zamora-Marín, Francisco J. Oliva-Paterna, Antonio Zamora-López & José F. Calvo

